# A *Rickettsiella* endosymbiont is a potential source of essential B-vitamins for the poultry red mite, *Dermanyssus gallinae*

**DOI:** 10.1101/2021.04.14.439672

**Authors:** Daniel R. G. Price, Kathryn Bartley, Damer P. Blake, Eleanor Karp-Tatham, Francesca Nunn, Stewart T. G. Burgess, Alasdair J. Nisbet

**Affiliations:** Moredun Research Institute, Pentlands Science Park, Edinburgh EH26 0PZ, United Kingdom; Pathobiology and Population Sciences, Royal Veterinary College, North Mymms, UK

**Keywords:** Endosymbiont, mutualist, symbiosis, Gammaproteobacteria, vitamin biosynthesis

## Abstract

Obligate blood-sucking arthropods rely on symbiotic bacteria to provision essential B vitamins that are either missing or at sub-optimal amounts in their nutritionally challenging blood diet. The poultry red mite *Dermanyssus gallinae*, an obligate blood-feeding ectoparasite, is primarily associated with poultry and a serious threat to the hen egg industry. Thus far, the identity and biological role of nutrient provisioning bacterial mutualists from *D. gallinae* are little understood. Here, we demonstrate that a *Rickettsiella* Gammaproteobacteria in maternally transmitted in *D. gallinae* and universally present in *D. gallinae* mites collected at different sites throughout Europe. In addition, we report the genome sequence of uncultivable endosymbiont “*Candidatus* Rickettsiella rubrum” from *D. gallinae* eggs. The endosymbiont has a circular 1. 89 Mbp genome that encodes 1973 protein. Phylogenetic analysis confirms the placement *R. rubrum* within the *Rickettsiella* genus, closely related to a facultative endosymbiont from the pea aphid and *Coxiella*-like endosymbionts from blood feeding ticks. Analysis of the *R. rubrum* genome reveals many protein-coding sequences are either pseudogenized or lost, but *R. rubrum* has retained several B vitamin biosynthesis pathways, confirming the importance of these pathways in evolution of its nutritional symbiosis with *D. gallinae. In silico* metabolic pathway reconstruction revealed that *R. rubrum* is unable to synthesise protein amino acids and therefore these nutrients are likely provisioned by the host. In contrast *R. rubrum* retains biosynthetic pathways for B vitamins: thiamine (vitamin B1) via the salvage pathway; riboflavin (vitamin B2) and pyridoxine (vitamin B6) and the cofactors: flavin adenine dinucleotide (FAD) and coenzyme A (CoA) that likely provision these nutrients to the host. We propose that bacterial symbionts which are essential to blood-feeding arthropod survival provide attractive targets for the development of novel control methods.

## Introduction

Animals live in a diverse bacterial world and mutualistic associations with bacteria can provide these animals with novel biochemical traits to exploit an otherwise inaccessible ecological niche (1). For example, specialist phloem-feeding insects of the order Hemiptera depend on bacterial endosymbionts to synthesise and provide essential amino acids that are largely absent in their phloem sap diet (2). Similarly, obligate blood-feeding arthropods, including insects, ticks and mites associate with nutritional mutualists that provide essential vitamins and cofactors that are in limited supply from their blood diet [recently reviewed in (3)]. Typically, the microbiome of obligate blood-feeding invertebrates is dominated by a single B vitamin provisioning symbiont. For example, the obligate blood-feeding African soft tick (*Ornithodoros moubat*) is associated with a *Francisella* (strain F-Om) mutualist that provides the host with essential B vitamins to supplement its blood meal diet (4). The genome sequence of *Francisella* F-Om bears the hallmarks of a typical host-restricted bacterial endosymbiont, with dramatic genome reduction resulting from loss of redundant genes that are not required for a symbiotic function. Importantly, *Francisella* F-Om retains biosynthesis pathways for B vitamins biotin (B7), riboflavin (B2), folic acid (B9) and cofactors coenzyme A (CoA) and flavin adenine dinucleotide (FAD) to supplement deficiencies in the hosts diet (4). This pattern of genome reduction and retention of B vitamin biosynthesis pathways is also observed in *Coxiella*-like endosymbionts (CLEs) from obligate blood-feeding ticks. Recent genome sequence studies revealed that, in comparison to the non-symbiotic pathogen *C. burnetii* (genome size 2.03 Mbp), CLEs from ticks have reduced genomes, as small as 0.66 Mbp for CLE from the lone star tick (CLE of *Amblyomma americanum*), yet they retain pathways for B vitamin and cofactor biosynthesis to supplement the nutritional requirements of their blood feeding host (5).

The poultry red mite (*Dermanyssus gallinae*) is an obligate blood feeder and a serious threat to the hen egg industry. Throughout Europe, *D. gallinae* prevalence is high, with up to 83% of commercial egg-laying facilities infested (6). Heavy infestations can reach up to 500,000 mites per bird and cause serious welfare issues, including anaemia, irritation and even death of hens by exsanguination (7). To utilise a blood meal as a single food source, our current hypothesis is that *D. gallinae* associates with nutritional mutualists which synthesize and supply essential B vitamins and cofactors that are absent in a blood diet. Previous studies have revealed that *D. gallinae* has a simple microbiome with 10 operational taxonomic units (OTUs) accounting for between 90% -99% of the observed microbial diversity (8). Here we identify a new species of Gammaproteobacteria from the *D. gallinae* microbiome, which we name “*Candidatus* Rickettsiella rubrum” sp. nov., which is vertically transmitted in *D. gallinae* and has reached fixation in European *D. gallinae* populations. Genome sequence analysis of *R. rubrum* reveals a moderately reduced genome of 1.89 Mbp with conserved biosynthesis pathways for B vitamins including thiamine (vitamin B1), riboflavin (B2), pyridoxine (B6) and the cofactors flavin adenine dinucleotide (FAD) and coenzyme A (CoA). Thus, *Rickettsiella rubrum* may synthesize and supply *D. gallinae* with essential nutrients that are missing in its blood diet.

## Methods

### Mite collection and endosymbiont-enriched DNA preparation

*Dermanyssus gallinae* were collected from a single commercial laying hen facility in the Scottish Borders, UK and maintained in 75 cm^2^ canted tissue culture flasks (Corning Inc., Corning, NY, USA) at 4°C for up to 4 weeks after collection. For experiments requiring mite eggs, freshly collected mixed stage and gender mites were placed into vented 25 ml Sterilin universal tubes and maintained at 25 °C, 75% relative humidity in a Sanyo MLR-350H incubator and eggs were collected the following day.

Since obligate bacterial endosymbionts are uncultivable outside the host, bacteria were derived from *D. gallinae* tissue lysates and host cells depleted using host depletion solution (Zymo Research, Irvine, CA, USA). Briefly, live mixed life-stage mites were surface sterilised with 70 % (v/v) ethanol for 30 s at room temperature followed by three 1 min washes in nuclease-free water. Mites (approx. 25 mg) were then homogenised in 200 µl nuclease-free water using a tube pestle and host cells lysed by addition of 1 ml of host depletion solution (Zymo Research, Irvine, CA, USA) with a 15 min incubation at room temperature with end over end mixing. Intact bacterial cells were pelleted by centrifugation at 10,000 x *g* for 5 min at room temp and DNA extracted from the pellet using a DNeasy® Blood & Tissue kit (Qiagen, Hilden, Germany). DNA concentration was assessed by the QubitTM dsDNA BR Assay Kit (Thermo Fisher Scientific, Waltham, MA, USA) and 1% (w/v) agarose gel electrophoresis.

### 16S rRNA amplicon sequencing and classification

Poultry red mite eggs were collected as described above and surface sterilised by two 5 min washes in 0.1% (w/v) benzalkonium chloride followed by two 5 min washes in 70% (v/v) ethanol. DNA was extracted from eggs using a DNeasy® Blood & Tissue kit (Qiagen, Hilden, Germany) with a lysozyme pre-treatment to lyse bacterial cells. DNA was quantified using a NanoDropTM One spectrophotometer (Thermo Fisher Scientific, Waltham, MA, USA) and DNA molecular weight determined on a 1% (w/v) agarose/TAE gel. A reagent-only control DNA extraction was performed in parallel using the same DNA extraction kit.

The presence of bacterial DNA in mite eggs was verified by PCR using universal bacterial 16S rRNA gene primers 27F-short (5’-GAGTTTGATCCTGGCTCA -3’) and 1507R (5’-TACCTTGTTACGACTTCACCCCAG -3’). Each 50 µl PCR reaction contained template DNA (100 ng), 1 U PlatinumTM *Taq* DNA Polymerase (Thermo Fisher Scientific, Waltham, MA, USA), 1x PCR buffer, 1.5 mM MgCl2, 0.2 mM of each dNTP and each primer at 0.2 µM. Cycling conditions were as follows: 94 °C for 2 min; 30 cycles of 94 °C 30 s, 58 °C 30 s, 72 °C 1min 30 s and a final hold of 72 °C for 10 min. A control PCR reaction was performed using the same conditions with an equivalent volume of eluate from the reagent-only control extraction. PCR products were cloned into pJET1.2 using the CloneJet PCR cloning kit (Thermo Fisher Scientific, Waltham, MA, USA) and transformed into chemically competent JM109 *E. coli* cells (Promega, Madison, WI, USA). Transformants were selected on Lysogeny broth (LB) agar plates containing 100 µg/ml ampicillin at 37 °C. Colony PCR was performed on randomly selected individual colonies using pJET1.2-F (5’-CGACTCACTATAGGGAGAGCGGC -3’) and pJET1.2-R (5’-AAGAACATCGATTTTCCATGGCAG -3’) vector primers using the previously detailed cycling conditions, except the primer annealing temperature was reduced to 56 °C. PCR products were analysed on a 1% (w/v) agarose/TAE gel and colonies containing the expected size amplification product were grown overnight in 10 ml LB containing 100 µg/ml ampicillin at 37 °C with shaking at 200 rpm. Plasmid DNA was isolated from each clone using Wizard® *Plus* SV Miniprep kit (Promega, Madison, WI, USA) and a total of 72 individual clones were sequenced with pJET1.2-F and pJET1.2-R primers at Eurofins Genomics Germany GmbH.

To assess the geographical association between *D. gallinae* and *Rickettsiella* we used DNA from a previously published mite collection from 63 sites across Europe (9). DNA from each collection sample was screened for *Rickettsiella* DNA using taxa specific 16S rRNA primers Rick-F (5’-GTCGAACGGCAGCACGGTAAAGACT -3’) and Rick-R (5’-TCGGTTACCTTTCTTCCCCACCTAA -3’), which were designed based on alignments in the PhylOPDb database (10). Each 25 µl PCR reaction contained template DNA (5 ng), 0.5 U PhusionTM High-Fidelity DNA Polymerase (Thermo Fisher Scientific, Waltham, MA, USA), 1x PCR buffer, 0.2 mM of each dNTP and each primer at 0.5 µM. Cycling conditions were as follows: 98 °C for 30 s; 30 cycles of 98 °C 10 s, 68 °C 30 s, 72 °C 30 s and a final hold of 72 °C for 10 min. PCR products were sequenced in both directions using Rick-seq-F (5’-AACGGCAGCACGGTAAAGAC -3’) and Rick-seq-R (5’-AGTGCTTTACAACCCGAAGG -3’) sequencing primers at Eurofins Genomics Germany GmbH.

16S rRNA sequences were classified with the RDP Classifier 2.13 (training set No. 18) (11) and sequences with <80% bootstrap support as their genus assignment were removed from the dataset. All remaining sequences were used in blastn searches against the GenBank database to identify their top hit.

### Genome Sequencing and Assembly

Raw PacBio reads generated from DNA isolated from *D. gallinae* eggs (published in (12)) were retrieved; the data set contained 7,318,092 reads for a total of 63,984,748,667 bases. Raw reads were mapped against the *D. gallinae* reference genome using Minimap2 v.2.17 (13) and unmapped reads were extracted from the resulting BAM files using SAMtools v1.11 (14). Unmapped reads (814,785 reads for a total of 1,274,422,647 bases) were assembled using the metaFlye assembler v.2.8.2 under default settings using the --pacbio-raw and --meta flags (15). The assembly containing 652 contigs was visualised with Bandage (16) which allowed identification of a circular 1.89 Mbp *Rickettsiella* genome with 12x coverage.

For massive parallel sequencing (MPS) host-depleted gDNA extracted as described above, was fragmented using a Covaris system, size-selected for 200 – 400 bp fragments and used for construction of a single strand DNA circle library. The library was amplified using phi29 DNA polymerase by rolling circle amplification to make DNA nanoballs (DNBs) and sequenced on a DNBSEQ-G50 platform as 150 bp paired end reads. Library construction and sequencing were performed by BGI Genomics (Shenzhen, China). This sequencing effort resulted in generation of 174,890,018 reads for a total of 26,233,502,700 bases. The reads were used to polish the *Rickettsiella* consensus sequence. Briefly, short-reads were mapped to the *Rickettsiella* genome using BWA-MEM aligner v0.7.17 (17) and base calls were corrected using five iterative rounds of polishing with Pilon v1.23 (18). The resultant assembly consisted of a single circular chromosome of 1,888,715 bp with 3,712x coverage.

### Genome Annotation

The genome was annotated using Prokka v.1.14.6 (19) and the automated pipeline included coding region prediction by Prodigal (20) and annotation of non-coding rRNAs using Barrnap and tRNAs using ARAGORN (21). As part of the Prokka pipeline, insertion sequences (IS) were annotated using the ISfinder database (22). Metabolic pathways for amino acids, B vitamins and cofactors were manually constructed using KEGG (23) and MetaCyc (24) reference pathways as guides and decorated with results from the genome annotation. The absence of genes in pathways was verified by tblastn searches against the *Rickettsiella* genome. The genome plot was generated using DNAplotter (25).

### Proposal for the species name of Rickettsiella-like endosymbiont

We have demonstrated that the symbiont belongs to the genus *Rickettsiella* and has less than 98.7% 16S rRNA sequence identity to its closest named phylogenetic neighbour, suggesting that the recovered genome is a new species. We propose in accordance with the terms for species designation for unculturable bacteria the name “*Candidatus* Rickettsiella rubrum” sp. nov. (hereafter *Rickettsiella rubrum* for simplicity). The specific name “rubrum” refers to the “red” colour of its mite host, the poultry red mite *Dermanyssus gallinae* after ingestion of a blood meal.

### Phylogenetic analysis

For phylogenetic analysis, full-length 16S rRNA sequences were aligned using ClustalW and a maximum-likelihood (ML) phylogenetic tree constructed using the Kimura 2-parameter (K2) model with gamma distributed with invariant sites (G+I). The substitution model was selected based on BIC score (Bayesian Information Criterion) and reliability of the tree was tested using bootstrap analysis (1000 replicates) with bootstrap values indicated on the tree. All phylogenetic analyses were performed using MEGA version X (26).

## Results and Discussion

### *Rickettsiella* is maternally inherited in *D. gallinae*

16S rRNA amplicon sequencing of DNA isolated from surface-sterilised *D. gallinae* eggs reveals that *Rickettsiella* is detectable in eggs (Figure 1), raising the possibility that *Rickettsiella* is maternally inherited in *D. gallinae*. It is notable that the *R. rubrum* whole genome sequence reported here was assembled from PacBio long-reads that were generated from DNA isolated from surface-sterilised mite eggs. In addition, a recent study of the *D. gallinae* microbiome identified *Rickettsiella* in all life-stages, including eggs, from mites collected from four geographically isolated commercial laying hen facilities in Czechia (27). Further attempts were made to assess the *Rickettsiella* transmission rate by running diagnostic PCR on DNA isolated from individual *D. gallinae* eggs, however, due to the small egg size and low recovery of DNA it was not possible to assess presence/absence in individual eggs.

**Figure 1.**
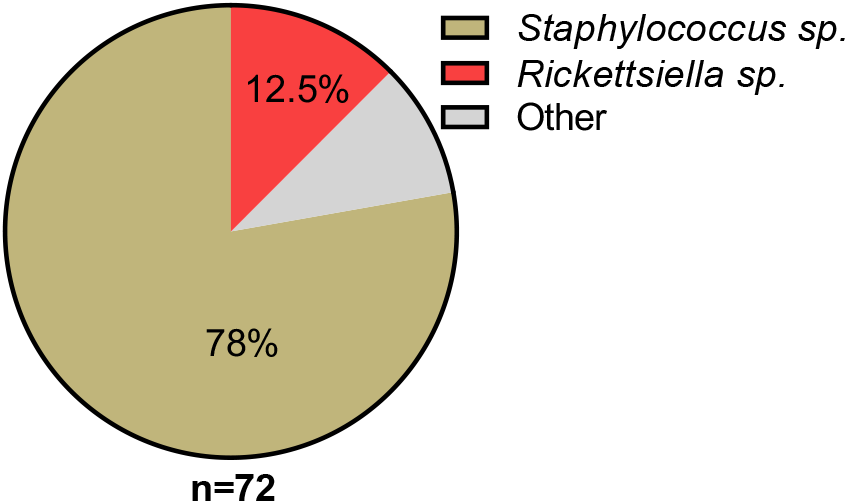
Classification and relative abundance of bacteria associated with *D. gallinae* eggs. The presence of bacterial DNA in mite eggs was verified by PCR using universal bacterial 16S rRNA gene primers. Amplicons were sequenced (n = 72) and classified with the RDP Classifier 2.13 (training set No. 18). Sequences with <80% bootstrap support as their genus assignment were removed from the dataset. Classifications are as indicated in the legend, other (grey) represents single hits (n = 1) to the following genera: *Blautia*; *Clostridium XII*; *Devosia*; *Paenalcaligenes*; *Salinicoccus*; *Streptococcus* and *Tsukamurella*.

### *Rickettsiella* infection has reached fixation in European populations of *D. gallinae*

We performed an extensive diagnostic PCR screen to test *D. gallinae* populations from collection sites throughout Europe for the presence of *Rickettsiella*. To do this, we used a previously prepared *D. gallinae* DNA collection extracted from mites sourced from commercial laying hen facilities from 62 sites across 15 European countries (9). For each sample site, total *D. gallinae* DNA was isolated from an individual adult mites, according to (9) and each sample was screened by diagnostic PCR using *Rickettsiella*-specific 16S rRNA primers. DNA samples from all *D. gallinae* sample sites (n = 62) were *Rickettsiella* positive, indicating that *Rickettsiella* infection has reached fixation in European *D. gallinae* populations (Figure 2).

**Figure 2.**
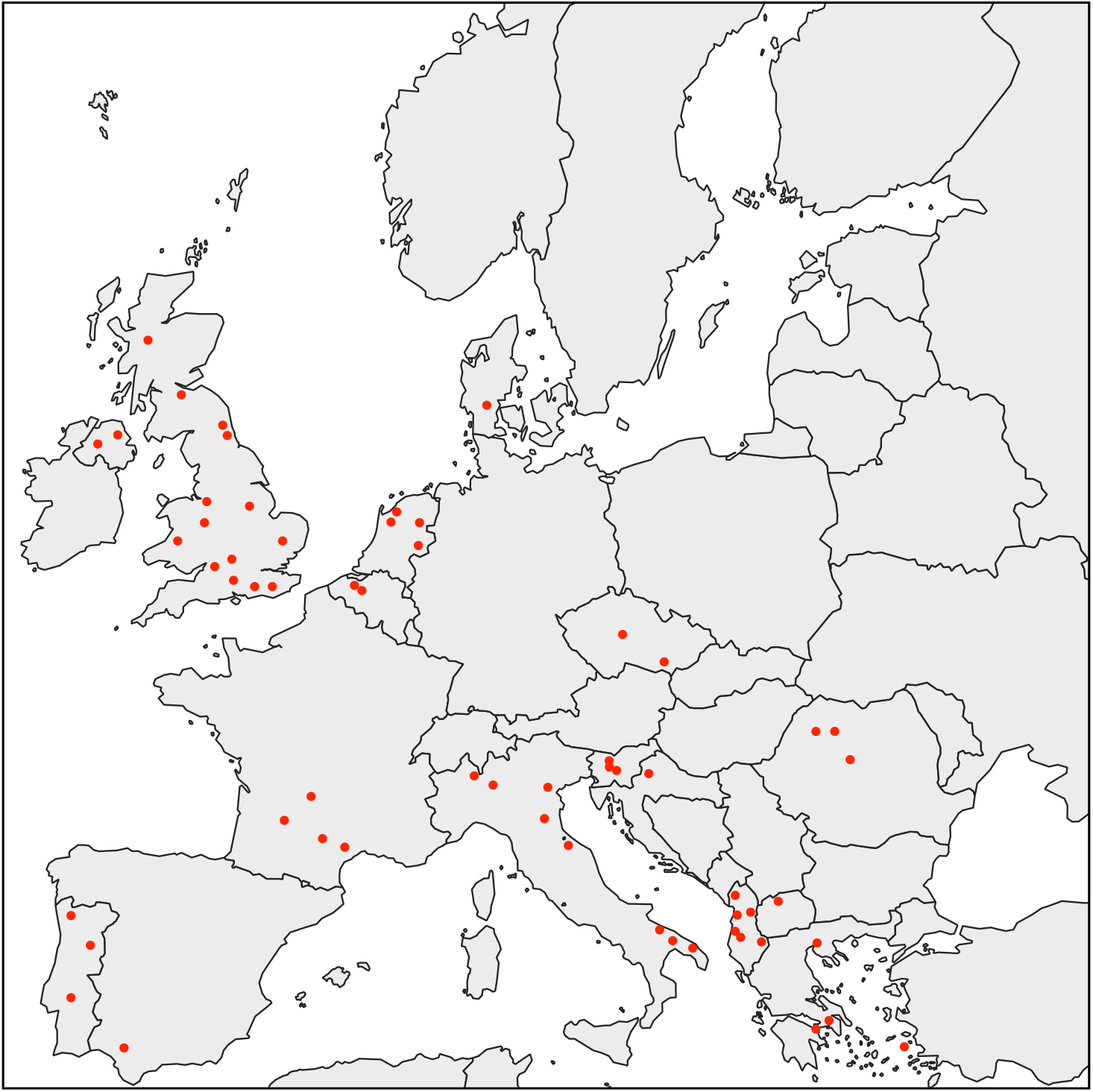
Map showing the distribution of *D. gallinae* populations analysed in this study. All individual adult female *D. gallinae* mites from each sampling site (63 sites across Europe) were positive for *Rickettsiella* infection (red circle) indicating *Rickettsiella* infection has reached fixation in European *D. gallinae* populations.

### General features of “*Ca*. Rickettsiella rubrum” genome

Previously generated PacBio long-read sequence data from *D. gallinae* eggs (12) were used to assemble the *R. rubrum* genome. From a total of 64.0 Gbp of sequence data, 1.3 Gbp of reads did not map to the *D. gallinae* draft genome and were used for metagenome assembly, resulting in generation of 652 contigs, which, after assembly, contained a circular *R. rubrum* chromosome of 1.89 Mbp. To correct errors associated with long-read sequence data, the *R. rubrum* assembly was polished using five iterative rounds of Pilon with DNBSEQTM short-read sequence data from symbiont enriched DNA. This yielded a circular chromosome of 1,888,715 bp with 3,712x coverage and a G+C content of 39.6 % (Figure 3). Based on Prokka gene prediction and annotation, the *R. rubrum* genome has 1,973 protein coding open reading frames (ORFs) with an average size of 870 bp which covered 91 % of the genome (Table 1 and Supplementary Table 1). Of these ORFs, 970 were assigned a biological function by Prokka annotation, 585 were annotated by BLAST homology to characterised proteins, while 227 matched hypothetical proteins of unknown function and 191 were unique to *R. rubrum*. In seven cases, pairs of adjacent genes were annotated with identical names and clearly the ORF was interrupted by a stop codon splitting the gene into two or more parts (these genes are highlighted in Supplementary Table 1). It is likely that these fragmented genes are non-functional and in the early stages of pseudogenization. To identify other pseudogene candidates we compared the length ratios of each predicted *R. rubrum* protein against their top blastp hit from searches against the NCBI nr protein database and flagged *R. rubrum* proteins that deviated by more than +/-25% (Supplementary Table 1). In summary, out of a total of 1,973 *R. rubrum* protein coding ORFs searched, only 312 (15.8%) deviate by more than +/-25% from their top hit and are candidate pseudogenes. However, it should be noted that the majority of these pseudogene candidates are “hypothetical proteins” of unknown function and therefore await experimental validation as genuine loss of function pseudogenes. We detected 41 tRNA genes (which can translate all 61 amino acid codons), 6 rRNA gene operons and 19 insertion-sequence (IS) elements.

**Table 1.** General genomic features of *R. rubrum* and allied Gammaproteobacteria

|  | *R. rubrum* | *R. viridis* | *C. burnetii*<br>RSA 493 | *E. coli* K-12 |
| --- | --- | --- | --- | --- |
| **Genome size, Mbp** | 1.89 | 1.58 | 2.00 | 4.64 |
| **G+C %** | 39.6 | 39.3 | 42.7 | 50.8 |
| **Protein-coding genes** | 1973 | 1362 | 1798 | 4242 |
| **Number of COGs<sup>#</sup>** | 1322 | 1033 | 1293 | 3812 |
| **Coding density %** | 91.0 | 87.1 | 77.7 | 85.8 |
| **Average gene size** | 870 | 1010 | 862 | 939 |

<sup>#</sup>COG, Cluster of Orthologous Groups

**Figure 3.**
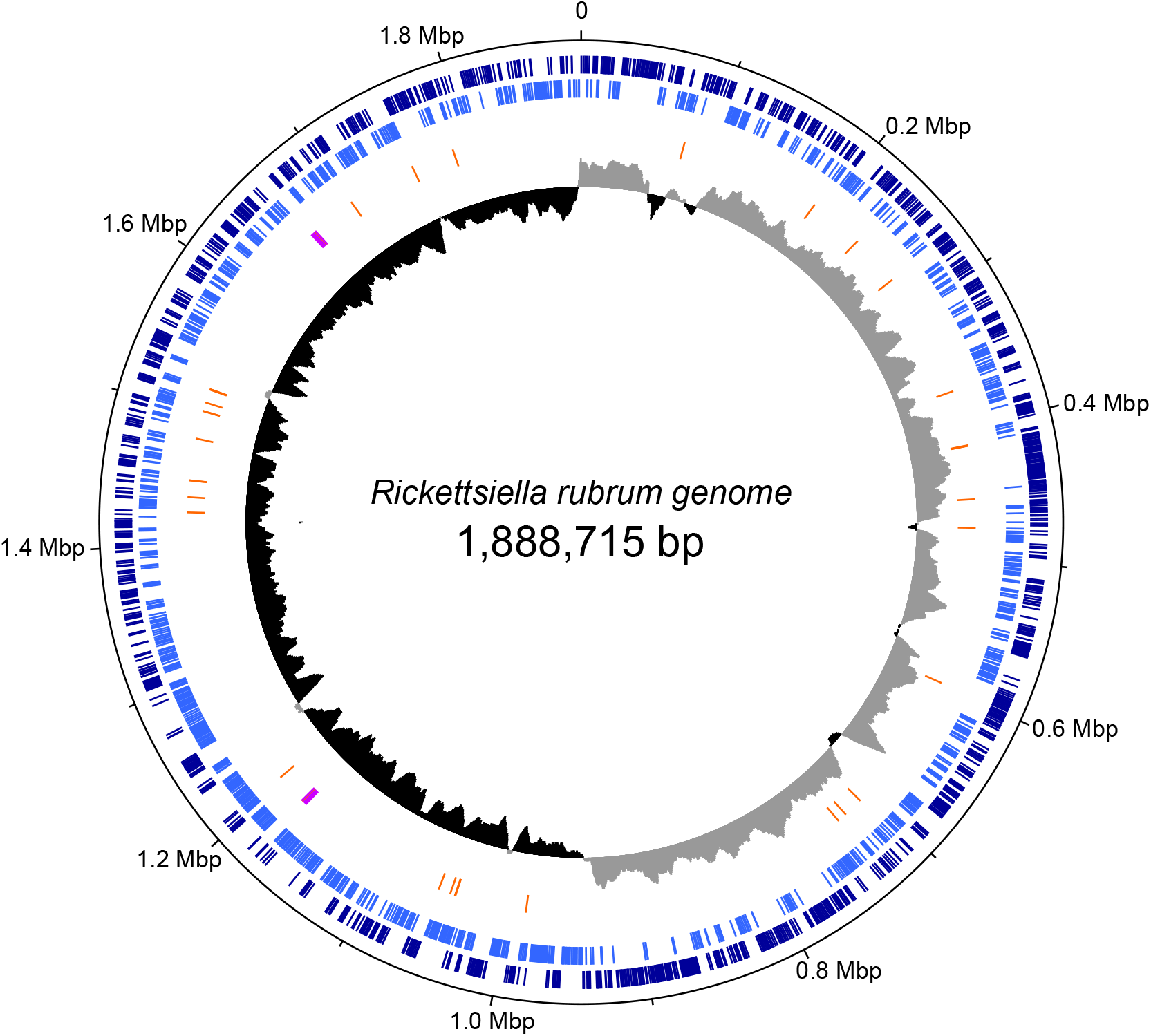
Map of the circular chromosome of “ca. Rickettsiella rubrum”. The innermost circle shows GC skew (window size: 10,000 bp) with grey and black indicating high (>0) and low (<0) (G-C)/(G+C) values. The second circle shows the positions of tRNA genes (orange) and rRNA genes (purple). The outer circles indicate the positions of protein coding genes on the plus strand (dark blue) and minus strand (light blue).

### *R. rubrum* is related to endosymbionts and endoparasites from the order *Legionellales*

All members of the Gammaproteobacteria order Legionellales are host-adapted for endosymbiotic or endoparasitic lifestyles within eukaryotic cells. Within Legionellales members of the genus *Rickettsiella* form a monophyletic group that diverged from *Coxiella burnetii*, the etiologic agent of Q fever, approx. 350 million years ago (29). *Rickettsiella* sp. are found in a wide range of arthropod hosts and are best known as obligate intracellular pathogens (29, 30), but recently some have been characterised as mutualistic endosymbionts (31, 32). Phylogenetic analysis, using 16S rRNA gene sequences from representative Gammaproteobacteria, confirms the placement *R. rubrum* within the *Rickettsiella* genus, closely related to the facultative endosymbiont *R. viridis* from the pea aphid *Acyrthosiphon pisum* (33) (Figure 4). In aphids, *R. viridis* was isolated from natural aphid populations and infection is associated with production of blue-green pigment molecules that accumulate in the host (32). Of particular note, aphids infected with *R. rividis* are not associated with negative impacts on host fitness and in some aphid strains infection is associated with elevated growth rates (32). Whole genome alignments between *R. rubrum* and *R. viridis* confirms that these two bacteria are very closely related but major genomic rearrangements including inversions, translocations and insertions are apparent (Figure 5).

**Figure 4.**
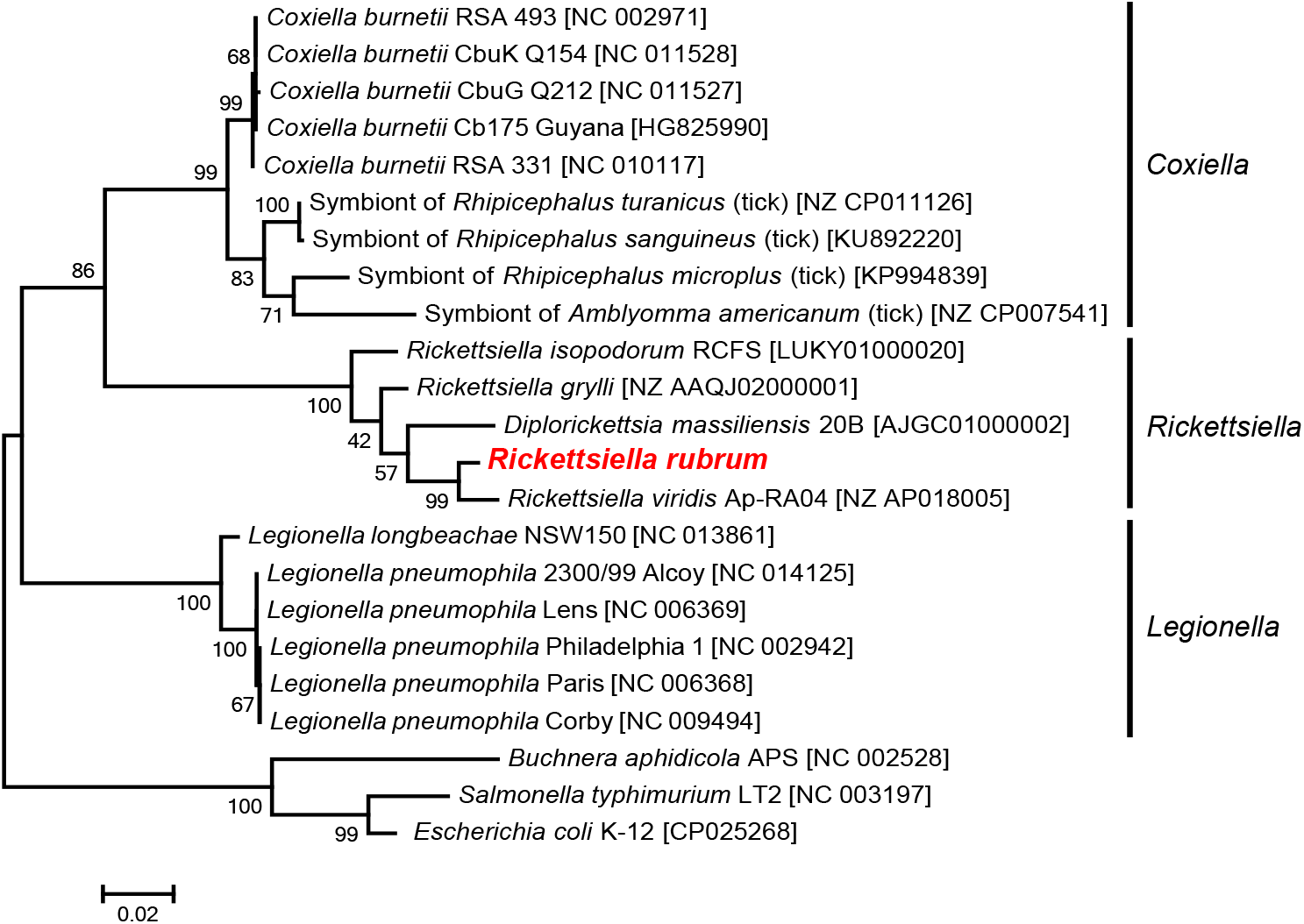
Phylogenetic placement of *R. rubrum* in the Gammaproteobacteria. The maximum likelihood phylogeny is inferred from 16S rDNA sequences. Statistical support is shown at each node from 1,000 bootstrap replicates. Scale bar represents 0.02 substitutions per site.

**Figure 5.**
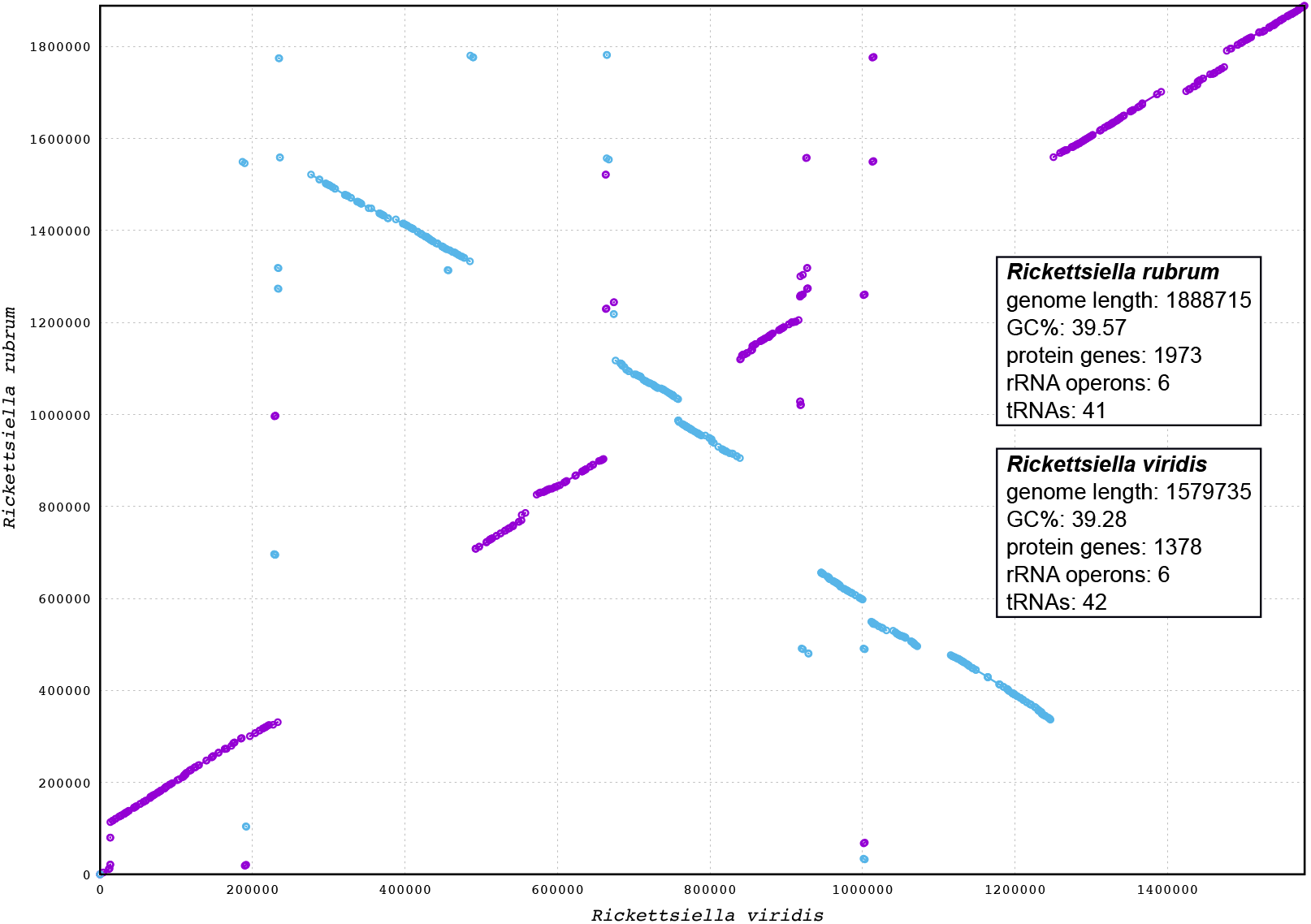
Synteny analysis between *R. rubrum* and *R. viridis* genomes. The *R. rubrum* genome is represented on the *y* axis and the *R. viridis* genome is represented on the *x* axis. Blue and purple lines represent synteny between the two genomes, with blue lines being inverted in *R. rubrum* relative to *R. viridis*.

Nutritional endosymbionts of blood-feeding arthropods are abundant in the order Legionellales. Again, closely related to *R. rubrum*, in the sister-genus *Coxiella* (Figure 4), *Coxiella*-like endosymbionts (CLEs) are required by ticks for supplementation of B vitamins that are absent in their blood meal and are essential for tick survival (34). In common with other host-restricted nutritional endosymbionts of arthropods, CLEs have massively reduced genomes, retaining functionally non-redundant genes that are essential for the symbiosis. Recent genome sequencing studies unveiled that, in comparison to *C. burnetii* (genome size 2.03 Mbp), CLEs from ticks exhibit extreme genome reduction, with genomes ranging from 0.66 Mbp for *Coxiella* sp. strain CLEAA (CLE of *Amblyomma americanum*) (5) to 1.73 Mbp for *Coxiella* sp. strain CRt (CLE of *Rhipicephalus turanicus*) (35). Presumably the range of genome size among CLEs of blood-feeding ticks reflects an ongoing dynamic process of reductive genome evolution. Metabolic reconstruction of these reduced genomes reveals intact B vitamin biosynthesis pathways, required for biosynthesis and provision of these essential nutrients to the host tick (5, 35).

In addition to ticks and mites, the blood-feeding louse *Polyplax serrata* is associated with a vertically transmitted, host restricted, nutritional endosymbiont from the genus *Legionella* (36). In comparison to *Legionella pneumophila* (genome size 3.4 Mbp), the etiologic agent of Legionnaires’ disease, the recently identified endosymbiont *L. polyplacis* has a massively reduced genome (0.53 Mbp) and parallels the reductive genome evolution observed in CLEs of blood-feeding ticks. Again, in the background of massive genome reduction *L. polyplacis* retains B vitamin biosynthesis pathways required for biosynthesis and provision of these essential nutrients to the host insect (36).

### Genomic reduction in *R. rubrum*: an ongoing process?

Genome reduction is widespread in maternally inherited bacterial endosymbionts and is associated with loss of genes that are functionally redundant within the host, resulting in compact endosymbiont genomes containing a subset of genes relative to their free-living ancestor (37). In general, “ancient” host-restricted endosymbionts have massively reduced genomes, for example, the smallest known cellular genome of the insect endosymbiont *Nasuia deltocephalinicola* is a diminutive 112 kbp and encodes just 112 protein coding genes, with distinctive adaptations to its host (38). In contrast, relatively “recent” host-restricted endosymbionts have much larger genomes, with gene content more reflective of their closest free-living ancestor. Following transition to a host-restricted lifestyle the genome of the newly acquired endosymbiont is associated with a period of genome instability, that typically includes a large increase in mobile elements in the genome, chromosomal rearrangements mediated by recombination between mobile elements and an increased pseudogene frequency (39, 40).

The genome of *R. rubrum* (1.89 Mbp) is only moderately reduced in comparison to closely related *C. burnetii* (2.03 Mbp) (Table 1), although it should be noted that *C. burnetii* is already host-adapted as an obligate intracellular parasite and as such, compared to free-living bacteria it has a degenerate genome (41). Relative to *C. burnetii*, the CLEs from blood-feeding ticks have further reduced genomes, typical of reduced genomes observed in other obligate nutritional endosymbionts of other blood-feeding insects and ticks, where the retained genes contribute to synthesis of essential B vitamins that are limited in the blood diet of their host (36, 42, 43, 4, 44). Perhaps the most striking example of genome reduction, in the transition from a pathogen to a nutritional mutualist, is the loss of virulence associated secretion systems: In the pathogens *C. burnetii* and *L. pneumophila* the type IV Dot/Icm secretion system (T4SS) functions to export a suite of virulence factors that modulate host physiology and are essential for establishment and maintenance of infection (41, 45, 46). Intriguingly, the massively reduced genomes of CLEAA and Ca. Legionella polyplacis from the blood feeding louse *Polyplax serrata* do not encode a Dot/Icm type IVB secretion system and presumably this secretion apparatus is not required in these nutritional mutualists (36, 5). In contrast, components of the Dot/Icm type IVB secretion system are retained in *R. rubrum* and are present in the closely related genomes of *R. viridis* and *R. gyrilli*, although the sequences of core components are highly divergent when compared with *C. burnetii* orthologs (Figure 6). It therefore remains to be determined if the Dot/Icm type IVB secretion system is functional in *R. rubrum* and if so what role it plays in cellular interactions with the host.

**Figure 6.**
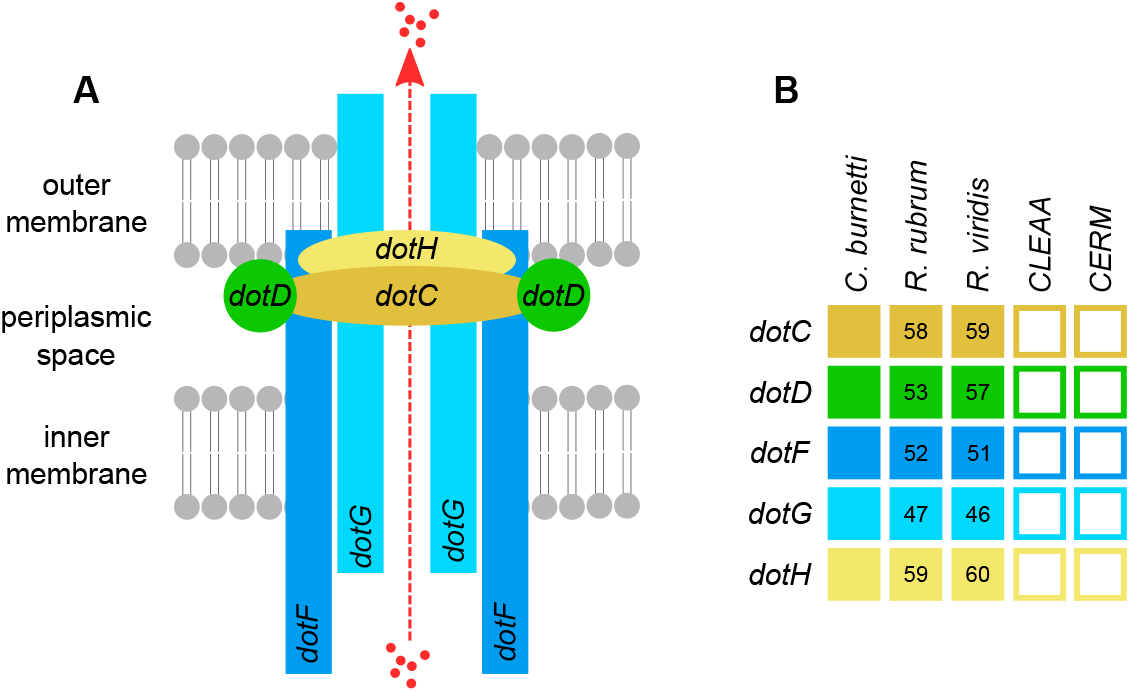
Comparative analysis of T4BSS (Dot/Icm) secretion system in *R. rubrum* and allied bacteria. **(A)** Representation of the core-complex of the Dot/Icm secretion system. **(B)** Presence (filled squares) and absence (white squares) of Dot/Icm components in genomes of *R. rubrum* and other species as indicated. Numbers in each square indicate percentage amino acid identity of each component relative to *C. burnetii*. CLEAA (*Coxiella* symbiont of *Amblyomma americanum*); CERM (*Coxiella* endosymbiont from *Rhipicephalus microplus*).

Genomes of other obligate intracellular bacteria typically have very few or no insertion (IS) elements, presumably due to the lack of opportunity for horizontal gene transfer (47, 48). In contrast, *R. rubrum* contains 19 IS elements evenly distributed around the genome and there are 8 copies of IS256 family transposase; 4 IS481; 4 ISNCY and 3 IS5. In addition, there is evidence of extensive horizontal gene transfer (HGT) within *R. rubrum* genome, including transfers from arthropods (2 HGT events), metazoa (6 HGT events), and numerous transfers from bacteria outside of Legionellales. Thus, the *R. rubrum* genome is highly dynamic as evident from the high number of HGT events, numerous IS elements and structural rearrangements in the *R. rubrum* genome relative to *R. viridis* (Figure 5). Based on these observations we conclude that *R. rubrum* is recently host-restricted with a genome of similar size to *C. burnetii* and is yet to undergo significant genome reduction as seen in other related blood-feeding CLE endosymbionts (5, 35).

### Metabolic capacity of *R. rubrum*: a putative nutritional mutualist

The *R. rubrum* genome, as with the related endocellular facultative symbiont *R. viridis*, retains genes for basic cellular processes including translation, replication, cell wall biosynthesis and energy production (Figure 7). In Supplementary Table S2, we provide a more detailed comparative gene content analysis between *R. rubrum* and genomes of *R. viridis* and *C. burnetii* using the pathway/gene list published by (38, 49). Surprisingly, both *R. rubrum* and *R. viridis* have a fragmented phospholipid biosynthesis pathway, suggesting that they are unable to complete *de novo* phospholipid biosynthesis. As phospholipid is an indispensable component of the cell membrane, phospholipid must either be imported from the host or the fragmented pathway is completed using host imported enzymes.

**Figure 7.**
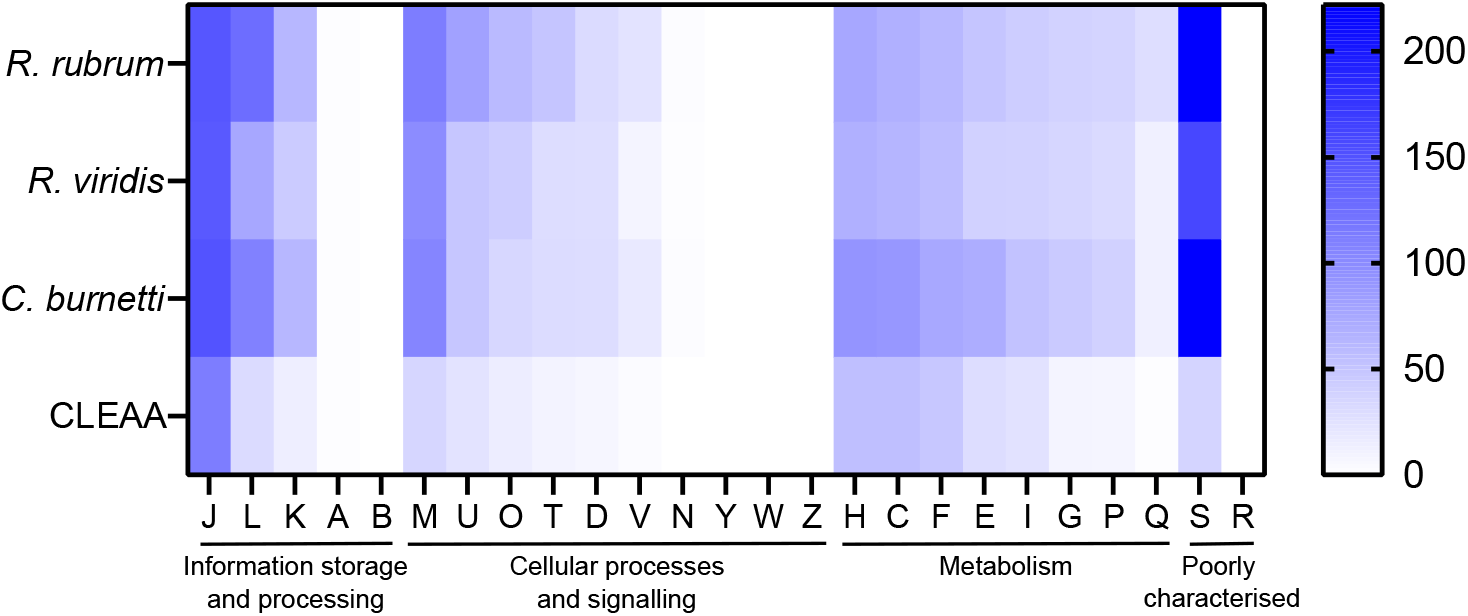
Heatmap comparison of Cluster of Orthologous Groups (COG) frequency in *R. rubrum* and related bacteria. Abbreviations for functional categories are as follows: **J**, Translation, ribosomal structure and biogenesis; **L**, Replication, recombination and repair; **K**, Transcription; **A**, RNA processing and modification; **B**, Chromatin structure and dynamics; **M**, Cell wall/membrane/envelope biogenesis; **U**, Intracellular trafficking, secretion, and vesicular transport; **T**, Signal transduction mechanisms; **O**, Posttranslational modification, protein turnover, chaperones; **D**, Cell cycle control, cell division, chromosome partitioning; **V**, Defense mechanisms; **N**, Cell motility; **Y**, Nuclear structure; **W**, Extracellular structures; **Z**, Cytoskeleton; **H**, Coenzyme transport and metabolism; **C**, Energy production and conversion; **F**, Nucleotide transport and metabolism; **E**, Amino acid transport and metabolism; **I**, Lipid transport and metabolism; **G**, Carbohydrate transport and metabolism; **P**, Inorganic ion transport and metabolism; **Q**, Secondary metabolites biosynthesis, transport and catabolism; **S**, Function unknown; **R**, General function prediction only. Scale bar (0, white; 200, blue) indicates number of COGs in each category.

Metabolic reconstruction of amino acid biosynthesis pathways revealed that *R. rubrum* is unable to synthesize protein amino acids and therefore these nutrients are likely provisioned by the host (Figure 8). The biosynthesis pathway for the essential amino acid arginine in mostly complete (8/9 required genes present), although precursor aspartic acid in not synthesized by *R. rubrum* and the bifunctional aspartokinase/homoserine dehydrogenase 1 (encoded by *thrA*) is missing, again suggesting this pathway is non-functional. Given that *D. gallinae* feeds on blood and is able to digest haemoglobin to release free amino acids (50), it likely has an excess of essential and non-essential amino acids that meet its own nitrogen requirements and those of *R. rubrum*. Indeed, in other nutritional endosymbnionts of obligate blood feeding arthropods, amino acid biosynthesis pathways are absent and it is likely the host supplies amino acids to the endosymbiont (5, 4, 45).

**Figure 8.**
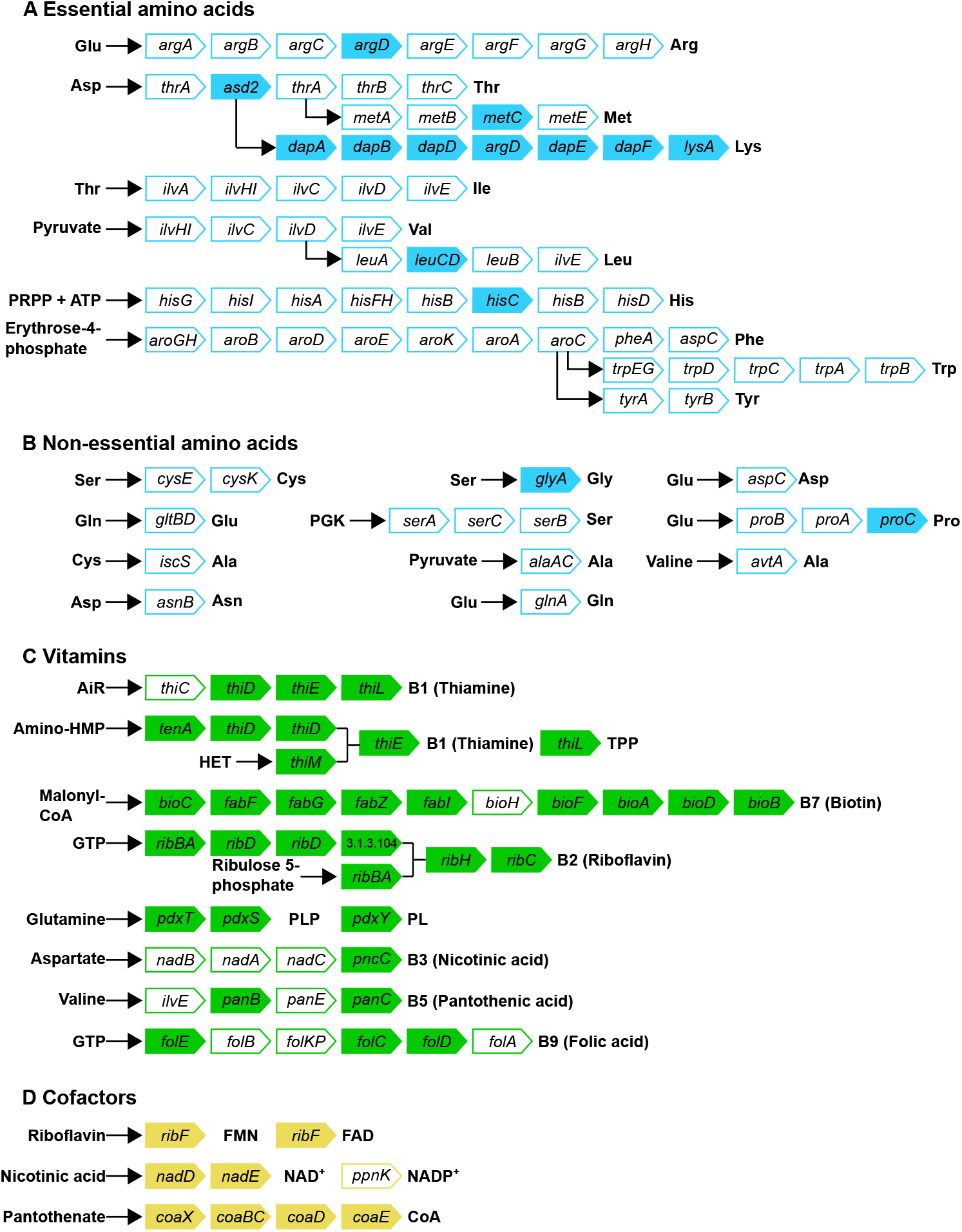
Biosynthetic pathways for synthesis of **A** essential amino acids; **B** non-essential amino acids; **C** vitamins and **D** cofactors in *R. rubrum*. Gene names are indicated in arrowed rectangles, coloured arrows show genes present in *R. rubrum*; missing genes are shown in white arrows.

Obligate blood feeding arthropods such as the human body louse (*Pediculus humanus*) (42), African soft tick (*Ornithodoros moubata*) (4) and the Lone star tick (*Amblyomma americanum*) (5) depend on nutritional endosymbionts to synthesize and provide B vitamins that are available in trace amounts in mammalian blood [reviewed in (3)]. To determine whether *R. rubrum* can play a similar role in *D. gallinae* we surveyed the *R. rubrum* genome for B vitamin biosynthesis genes. The *R. rubrum* genome has conserved genes involved in the biosynthesis of seven B vitamins, including complete biosynthetic pathways for thiamine (vitamin B1) via the salvage pathway, riboflavin (vitamin B2), pyridoxine (vitamin B6) and the cofactors flavin adenine dinucleotide (FAD) and coenzyme A (CoA) (Figure 8). The biosynthesis pathway for biotin (vitamin B7) is largely complete (9/10 genes present) although it is missing *bioH*, which is required for pimeloyl-CoA synthesis. The annotated biotin biosynthesis pathway is based on that of the model organism *E. coli*, where *bioC* and *bioH* are required for synthesis of the intermediate pimeloyl-CoA. However, unlike the representative “*bioC*/*bioH*” pathway of *E. coli* many *bioC*-containing microorganisms lack *bioH* homologues, raising the possibility of non-homologous gene replacement in some bacteria (51). To date, there are five documented cases of *bioH* gene replacement, which includes *bioK* of *Synechococcus* (51), *bioG* of *Haemophilus influenzae* (51), *bioJ* of *Francisella sp*. (52), *bioV* of *Helicobacter sp*. (53) and *bioZ* of *Agrobacterium tumefaciens* (54). Further tblastn searches against the *R. rubrum* genome using *bioH* and the non-homologous gene replacements *bioK, bioG, bioJ, bioV* did not identify gene products that can fill the *bioH* gap. However, a gene encoding ketoacyl-ACP synthase (KAS) III from *R. rubrum* has similarity to *bioZ* of *A. tumefaciens* and is therefore a candidate to replace *bioH*. Given the retention of a long biotin biosynthesis pathway in *R. rubrum* (9/10 genes present) and the propensity for the missing *bioH* gene to be replaced in other bacteria, we predict that the biotin biosynthesis pathway is functional in *R. rubrum*. In contrast, the other B vitamin biosynthesis pathways for nicotinic acid (vitamin B3), pantothenic acid (vitamin B5) and folic acid (vitamin B9) are more fragmented and thus may be non-functional. Although *R. rubrum* biosynthesis pathways for vitamin B3, B5 and B9 are fragmented future work will analyse these pathways in the context of the *R. rubrum* metagenome. Genome analyses of other nutritional endosymbionts reveal that some retained “broken” pathways are functional with gene products supplemented from multiple species of symbiont partners resulting in metabolic mosaics for the synthesis of essential nutrients (55, 56). We know from 16S rRNA amplicon sequencing that the *D. gallinae* microbiome is relatively simple, with 10 OTUs accounting for between 90% -99% of the microbial diversity observed (8). Currently, the contribution of other partners in the *D. gallinae* microbiome towards B vitamin biosynthesis is unknown and will be the target of future studies.

## Author Contributions

DRGP, AJN, STGB conceived the study. All authors designed the research. DRGP, EKT performed research. DRGP, AJN, STGB analysed data. DRGP wrote the paper with contributions from all authors. All authors read and approved the final manuscript.

## Data availability statement

DNBseq reads were deposited to the Sequence Read Archive (SRA), under NCBI BioProject PRJNAXXXXXX. The *R. rubrum* genome assembly is available under NCBI BioProject PRJNAXXXXXX.

## Acknowledgements

We thank the Bioservices Group at Moredun Research Institute for their ongoing help and expertise and UK farmers for allowing access to sites for *D. gallinae* collection.

